# Extremely fast and incredibly close: co-transcriptional splicing in budding yeast

**DOI:** 10.1101/083170

**Authors:** Edward W.J. Wallace, Jean D. Beggs

## Abstract

RNA splicing, an essential part of eukaryotic pre-messenger RNA processing, can be simultaneous with transcription by RNA polymerase II. Here, we compare and review independent next-generation sequencing methods that quantify co-transcriptional splicing in budding yeast. Splicing in yeast is fast, taking place within seconds of intron transcription, while polymerase is within a few dozens of nucleotides of the 3’ splice site. Ribosomal protein mRNAs are spliced particularly fast and co-transcriptionally. Intron-mediated regulation of some genes is also likely to be co-transcriptional. We suggest that intermediates of the splicing reaction, missing from current datasets, may hold key information about splicing kinetics.

**Trends:**

- Independent next-generation sequencing methods quantify co-transcriptional splicing in budding yeast
- Ribosomal protein mRNAs are spliced particularly fast and co-transcriptionally
- Intron-mediated regulation of *DBP2* and *RPS9A* is likely co-transcriptional
- Splicing intermediates, missing from current datasets, may hold key information

## Introduction

RNA splicing is an essential process in the maturation of most transcripts produced by eukaryotic RNA polymerase II (Pol II). RNA molecules can be spliced while still being transcribed, as shown by pioneering electron micrography studies of Drosophila embryo transcripts (Beyer et al., 1981, 1988). Now, there is extensive support for functional coupling between splicing and transcription (reviewed in Alexander & Beggs, 2010, Naftelberg et al., 2015, Saldi et al., 2016). Evidently, transcription can affect splicing and vice-versa, but how this is achieved and regulated is largely unknown. The speed of splicing varies from gene to gene, depending on the strength of splice sites as well as other factors, and expression of some genes is regulated through splicing. What are the transcriptome-wide patterns of splicing kinetics?

Recently, distinct next-generation sequencing approaches have measured the coupling of transcription to splicing in *Saccharomyces cerevisiae*: fast metabolic labelling with 4-thio-uracil (4tU-seq; Barrass et al., 2015), nascent RNA-seq (Harlen et al., 2016) and single molecule intron tracking (SMIT; Carrillo Osterreich et al., 2016) These approaches produce, for many genes, *quantitative* estimates of the speed of splicing, extent of co-transcriptional splicing, or polymerase position at splicing, respectively. Collectively, these data demonstrate that, in budding yeast, most introns are spliced out very soon after the intron is transcribed, a scenario that was controversial ten years ago. Alternative next-generation sequencing methods measure polymerase position, without so far providing high-resolution measures of splicing: nascent elongating transcript sequencing (NET-seq; Harlen et al., 2016), and crosslinking to modified Pol II and analysis of cDNA (mCRAC; Milligan et al., 2016). Importantly, Harlen et al. and Milligan et al also measure how the phosphorylation state of the carboxy-terminal domain (CTD) of Pol II large subunit correlates with splicing factor recruitment or intron position genome-wide. Here, we discuss the strengths and limitations of each approach, and the extent to which their results agree. One remarkable point of agreement is that ribosomal protein transcripts, the largest and most abundant class of spliced mRNAs, tend to be spliced faster and more co-transcriptionally.

## Methods of measuring nascent RNA

### 4tU-seq (Barrass et al., 2015)

4tU-seq uses short-timescale metabolic labelling to measure the first minutes of RNA transcription and processing. Nascent transcripts are labelled by incorporation of the uracil analogue 4tU. After flash-freezing and cell lysis, the 4tU-labelled transcripts are biotinylated and then isolated based on their affinity for streptavidin beads, followed by randomly primed reverse transcription and sequencing. Using a statistical model to compare the spliced:unspliced ratio of transcripts labelled with 4tU for different times, the *relative speed of splicing* of each transcript is determined. The reported measure is the area under the curve (AUC) of the spliced:unspliced ratio, a time-weighted average of the proportion spliced within 5 minutes of synthesis. Note that the overall rate of conversion of pre-mRNA to spliced mRNA is determined in real time rather than relative to the movement of Pol II.

As 4tU-seq is essentially RNA-seq applied to the subset of RNA that is newly synthesised, a range of mature, validated tools are available to analyse the data, including building on the MISO algorithm for mRNA isoform quantification (Katz et al., 2010), to accommodate time-series labeling (Huang & Sanguinetti, 2016). These statistical tools provide robust gene-level quantification of splicing by combining strong information from the small proportion of reads that span a splice junction with individually weak information from the majority of non-junction reads. However, 4tU-seq is uninformative about splicing intermediates (the products of the first catalytic step) or excised introns (the by-product of the second catalytic step): indeed, at these short timescales the intron:exon ratio may detect variability in degradation rates of excised introns as well as variability in splicing rates (box 1).

##### Box 1: Difficulties in detecting all stages of splicing

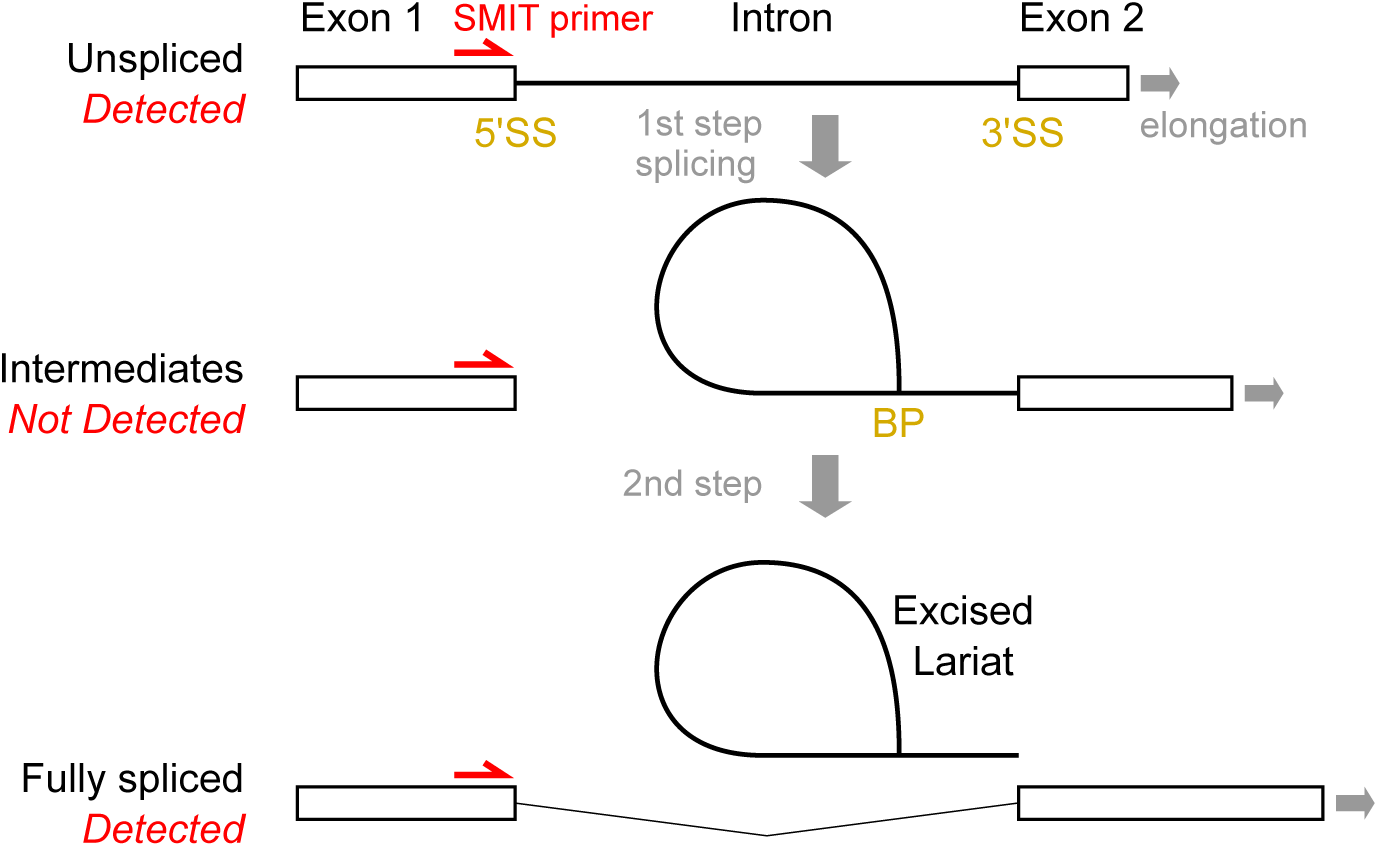

The pre-mRNA splicing reaction has 2 catalytic steps: in the first step, the 5’SS is cleaved and the lariat intron-exon intermediate species is formed by joining at the branch point (BP), and in the second, the 3’ end of the upstream exon is joined to the downstream exon at the 3’SS, resulting in spliced mRNA and an excised lariat intron. Biased detection of these various RNA species complicates analysis in all methods discussed here.

4tU-seq quantifies splicing by the ratio of exonic to all reads for a given RNA, as well as incorporating the unique information from junction reads, implicitly assuming that excised introns are degraded quickly enough that most intronic reads are from unspliced pre-mRNA. Although excised introns are, in general, quickly degraded, some lariats might be degraded on a timescale comparable to that of 4tU-seq measurements. Indeed, some evolutionarily conserved non-coding RNAs, including snoRNAs, are processed from introns, and these may be degraded slowly (Hooks et al 2016). Therefore, 4tU-seq may *under-estimate* the rate of splicing for transcripts with stable intron-derived products.

SMIT quantifies splicing by sequencing PCR products that extend from a primer site upstream of the 5’SS to the 3’ Pol II position. For splicing intermediates, these sites are not contiguous, and so are not detectable by SMIT. Therefore, SMIT may *over-estimate* the extent of splicing of transcripts for which lariat intermediates represent a significant proportion of the total (slow second step), and *over-estimate* the speed of splicing if spliceosome assembly and the first step of splicing occur before the 3’SS is transcribed (first step splicing is possible once the BP becomes available), such that distance from the 3’SS does not reflect all aspects of splicing.

Consider nascent transcripts whose 3’ end is at position *n*. If the true count of pre-mRNA is ***P*_*n*_**, splicing intermediates is ***I*_*n*_**, and spliced mRNA is ***F*_*n*_**, then the proportion that has completed splicing is 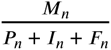. However, SMIT estimates the proportion spliced as 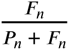, which is strictly greater; the difference depends on ***I*_*n*_**.

This may explain a puzzle in the SMIT analysis, that some estimates of the position of onset of splicing conflict with structural models. The *average* position for onset (10% of transcripts spliced) of 26 nts after transcription of the 3’ splice site is remarkably soon given that the distance between polymerase and spliceosome active sites is estimated as 24 nts (Carrillo Osterreich et al., 2016). Perhaps this is not surprising: in a first-order kinetics approximation of splicing, the relative time or position of splicing would be an exponential distribution, with a mode at the first available point. However, beyond averages, SMIT’s quantification algorithm reports that 24 of 87 measured genes have splicing onset less than 24nts after the 3’ SS, 9 are 50% spliced by then, and onset is not reported for a further 31 genes.

NET-seq detects Pol II-associated 3’ ends of RNA species, including many reads exactly at the 3’SS which are most likely from excised introns attached to spliceosomes that are still associated with Pol II (Harlen et al., 2016). Interestingly, an earlier NET-seq dataset observed an abundance of 3’ end reads exactly at the 5’SS (Churchman and Weissman 2011), also consistent with a population of splicing intermediates that pull down with elongating Pol II.

Complementary methods quantify splicing intermediates by sequencing, even in wild-type cells (Gould et al., 2016; Qin et al., 2016). However, these rely on enrichment of lariats, so provide weak information on the kinetics of splicing. Methods that measure all steps of the co-transcriptional splicing reaction simultaneously are thus needed to fill in the gaps in our quantitative understanding of co-transcriptional splicing.

### SMIT (Carrillo Osterreich, et al., 2016)

The SMIT approach cleverly exploits paired-end RNA sequencing to measure the splicing status of nascent transcripts that are assumed to be still associated with Pol II. Cells are harvested by centrifugation and washed in cold buffer, requiring the assumption that transcription and splicing are arrested to the same degree during sample preparation, then chromatin is isolated in a multi-step procedure developed earlier by the same group (Carrillo Osterreich, et al., 2010), and mature polyA+ RNA is further depleted. Remaining chromatin-derived RNA is reverse-transcribed from a 3’ end-ligated oligonucleotide, then PCR amplified, using gene-specific primers upstream of target introns to ensure high read-depth for target transcripts. Most budding yeast intron-containing genes have short first exons, therefore 87 target genes were selected based on the suitability of first exons to prime PCR across the intron. To aid correct quantification despite dense sampling of target genes, random molecular barcodes were incorporated in the ligated oligonucleotide. By design, the 3’ end reads report polymerase position, while the 5’ end reads, because they start close to the introns, report splicing status. SMIT detects intron-containing transcripts prior to the first step of splicing and also detects spliced transcripts that have completed the second step of splicing (exon joining, box 1). SMIT cannot detect transcripts that have undergone only the first step of splicing, for which the 3’ end of nascent RNA is not contiguous with the 5’ primer site (box 1). The proportion of transcripts that have undergone both steps is calculated at spatially binned Pol II positions for each tested gene, and the position is estimated at which 10%, 50%, and 90% of nascent transcripts are spliced comparably to “saturation”, i.e. splicing ratio for the most 3’ Pol II positions. Thus SMIT reports the *distribution of Pol II positions at splicing*. SMIT’s measurements of relative positions of polymerase and splicing are analogous to those made from the electron micrograph tracings of Beyer et al., (1981,1988).

Importantly, Carrillo Osterreich et al. validate the SMIT assay by comparison to other methods: long-read sequencing confirms the onset of splicing soon after polymerase passes the 3’SS. Further, SMIT shows that splicing moves further 3’ from the 3’SS in a fast-elongating Pol II mutant, as predicted by kinetic competition of polymerization and splicing.

### Nascent RNA-seq (Harlen et al., 1016)

For nascent RNA-seq, cells with affinity-tagged Pol II are flash-frozen and lysed, Pol II is immunoprecipitated and the associated, nascent RNA transcripts are purified, fragmented and sequenced. This approach reports what fraction of RNA is spliced co-transcriptionally, but does not provide information about Pol II occupancy or splicing kinetics. As for 4tU-seq, only a small proportion of nascent RNA-seq reads span a splice junction, thus for less abundant transcripts the fraction spliced co-transcriptionally is the ratio of two small numbers that are susceptible to count noise. Alternatively, one can compare the nascent RNA-seq coverage in a window upstream and downstream of the 3’SS.

### NET-seq (Churchman and Weissman, 2011; Harlen et al., 2016)

NET-seq quantifies Pol II occupancy genome-wide by sequencing the 3’ ends of all transcripts associated with Pol II (Churchman & Weissman, 2011). Similar to nascent RNA-seq, RNA is extracted from immunoprecipitated Pol II, the crucial difference being that short reads (75 nt) are obtained from the *3’ ends* of extracted RNA. As only a very small proportion of all nascent transcripts are spliced within these short distances, there is a very small number of spliced reads, so this approach is not suited to measuring splicing relative to Pol II position.

Detection of tRNA, which is produced by RNA pol III, and accumulation of reads at the 3’ ends of snoRNAs, suggests detectable levels of contamination from mature transcripts in NET-seq data. Similarly, 4tU-seq detects non-Pol II transcripts by design, and low levels of background RNA are isolated even without 4tU-labeling.

### mCRAC (Milligan et al., 2016)

With CRAC (Cross-Linking and Analysis of cDNA; Granneman et al., 2009), short fragments of RNA are identified that are UV crosslinked to, and protected from nuclease digestion by, a tagged bait protein. The protocol is analogous to CLIP (Darnell 2010), but features a stringent denaturation step to reduce background. Modification CRAC (mCRAC) employs a two-step purification, first with tagged Pol II, then with antibodies specific to different phosphorylation states of Pol II’s CTD, to obtain fragments of nascent transcripts associated with distinct states of the elongation machinery. Importantly, these data were used as input to a Bayesian model that suggests multiple distinct initiation and elongation states of Pol II. We address this dataset only briefly here on account of its relatively low depth of sequencing that does not permit an assessment of co-transcriptional splicing.

The data obtained by these approaches measure many aspects of transcription beyond those coupled to splicing: 4tU-seq, NET-seq and mCRAC allow estimates of transcription rate as distinct from RNA abundance, including of antisense and unstable transcripts, and a comparison would be instructive. However, we focus here on coupling of transcription and splicing.

## Results

### Ribosomal protein genes tend to be spliced fast and co-transcriptionally

There is a great deal of agreement between results of these distinct methods (Fig. 1A-C). Importantly, SMIT shows that the second step of splicing is complete on most transcripts when Pol II has moved less than 100 nt beyond the 3’SS, consistent with nascent RNA-seq’s measure that the majority of splicing is co-transcriptional (Fig. 1A). Also, transcripts that are seen to be spliced fast by 4tU-seq are generally measured as spliced co-transcriptionally by SMIT (Fig. 1B) and nascent RNA-seq (Fig. 1C): in particular, ribosomal protein (RP) transcripts. Transcripts measured by SMIT to be spliced when Pol II is further from the 3’SS appear to be spliced more slowly by 4tU-seq analysis, including the well-characterised *ACT1*. Furthermore, *YRA1*, which is negatively auto-regulated by an intron-dependent mechanism (Dong et al 2007; Preker & Guthrie, 2006), is spliced distally, slowly, and inefficiently (Fig. 1A,B).

**Figure 1:**
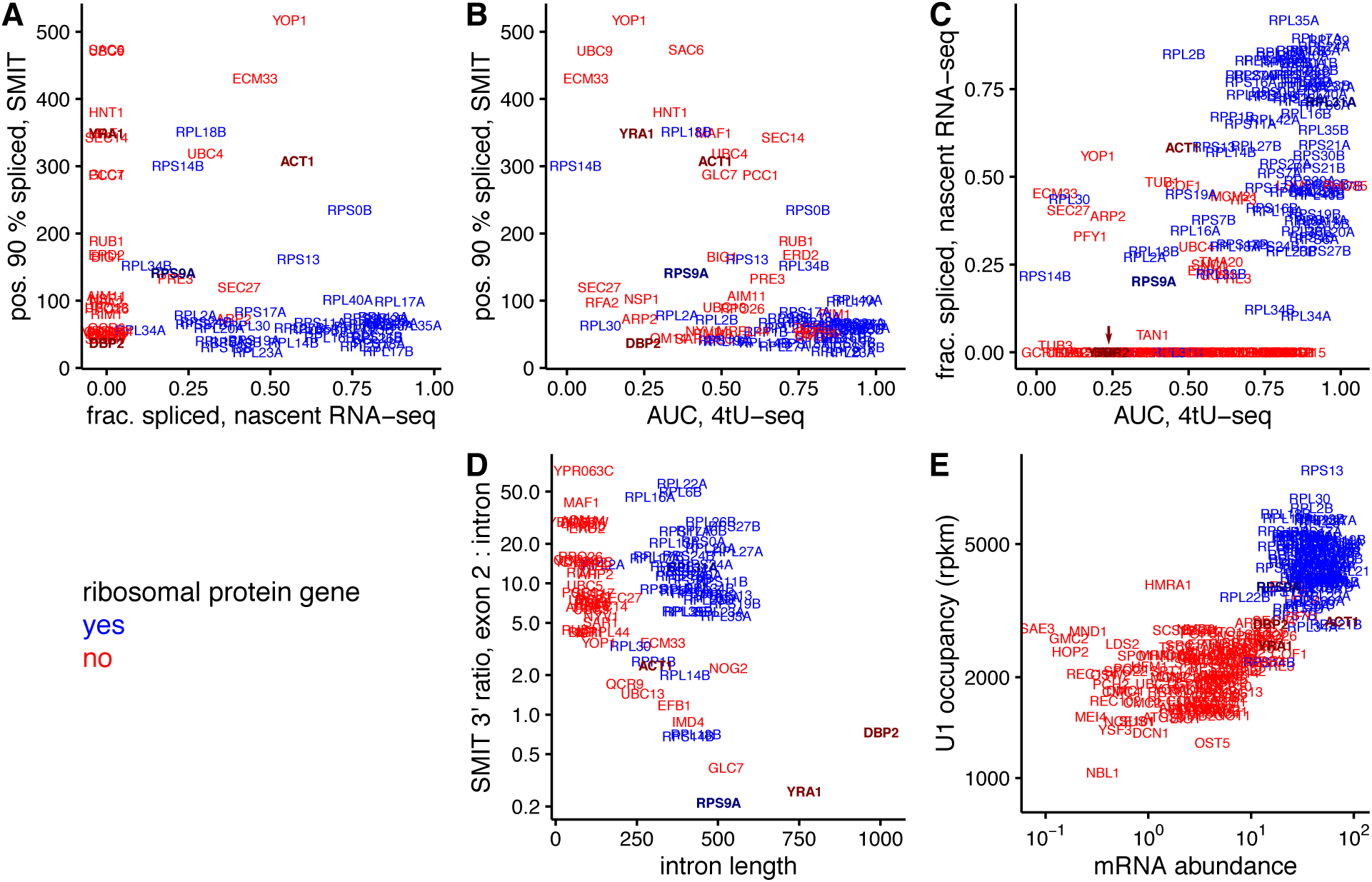
Intron-containing ribosomal protein transcripts (blue) tend to be spliced faster and more co-transcriptionally, compared to non-ribosomal transcripts (red). Gene names are plotted with centre indicating the position on each scale; select genes mentioned in text are shaded and bolded, and overlaid YRA1 and DBP2 in Fig. 1C indicated with an arrow. A. The distance in nts of Pol II from the 3’SS when 90% of transcripts are spliced according to SMIT (pos. 90% spliced; Carrillo-Osterreich et al., 2016) is plotted versus fraction spliced co-transcriptionally measured by nascent RNA-seq (Harlen et al., 2016). B. Pos. 90% spliced by SMIT versus AUC by 4tU-seq (a weighted average over the proportion spliced at 1.5, 2.5 and 5 minutes; Barrass et al 2015). C. Fraction spliced according to nascent RNA-seq versus AUC by 4tU-seq. D. Ratio of SMIT 3’ ends in introns and exon 2 (spliced or unspliced; Carrillo Osterreich et al., 2016) versus intron length. E. U1 occupancy measured by ChIP-nexus (Harlen et al., 2016) versus mRNA abundance (Csardi et al., 2015).

RP transcripts have other distinguishing patterns in these data: they have relatively fewer SMIT 3’ end reads in the intron compared to exon 2, despite their longer introns (Figs. 1D, S1). This suggests a large decrease in polymerase speed from intron to exon 2, consistent with NET-seq. Furthermore, RP genes have higher U1 occupancy as measured by ChIP-nexus (Harlen et al., 2016), so that the reported “high U1 occupancy genes” are essentially synonymous with the RP genes (Fig. 1E). Notably, the U1 occupancy differs only 3-fold between RP and non-RP genes, which is much less than the difference in mRNA abundance or, presumably, transcription rate. This is consistent with a low dynamic range or high background signal for U1 occupancy as measured by ChIP, but is also consistent with RP transcripts recruiting more U1 but for shorter times, due to their faster splicing.

These assays reveal differences in splicing between paralogous transcripts, for example of the *RPS14A* and *RPS14B* genes. It has been shown that excess S14 protein can bind to a stem-loop structure in *RPS14B* pre-mRNA, inhibiting its splicing and leading to its rapid degradation (Fewell and Woolford, 1999). The 4tU assay shows *RPS14B* transcripts splicing much more slowly than *RPS14A* transcripts; nascent RNA-seq shows almost twice the proportion of spliced reads for *RPS14A*. SMIT quantified *RPS14B* as spliced late compared to other RPs, but did not measure *RPS14A*.

### Apparent discrepancies between the assays

Curiously, some transcripts are apparently spliced slowly or less efficiently, yet close to the 3’SS (lower left quadrant in Fig. 1B). This apparent paradox could reflect the distinction between time and position of splicing: Pol II elongating more slowly or pausing near the 3’SS while splicing takes place would allow even slow splicing to occur co-transcriptionally. Artifacts in one or more of the assays could also explain the discrepancy: if an excised intron were degraded particularly slowly and sequenced efficiently, that would depress 4tU-seq estimates of splicing speed. Likewise, SMIT could overestimate the proportion of fully spliced transcripts near the 3’SS if splicing intermediates represent a large proportion of slowly spliced transcripts, as they are not detected in this assay (Box 1).

The transcripts that are apparently spliced most slowly (by 4tU-seq), and 3’SS proximally (by SMIT), are: *SEC27, RFA2, NSP1, ARP2, DBP2, OM14, SAR1, RPL2A, RPS9A*, and *RPL30* (Fig 1B). Notably, *DBP2, RPL30*, and *RPS9A* are all known to be negatively regulated by proteins binding to their introns, to which we return later. *RPL2A* is measured as proximally spliced by SMIT, thus may be a potential candidate for transcriptional pausing. *ARP2* and *OM14* are both spliced inefficiently and proximally, but with SMIT saturation values of only 59% and 20% respectively; in this respect the assays agree. The apparent discrepancy is caused by normalizing the low proportion of proximally-spliced reads by the still-low proportion of distally-spliced reads. For the others we have no alternative explanation.

### Intron-mediated inhibition may be co-transcriptional

The Rpl30 protein negatively regulates splicing of the *RPL30* transcript by binding co-transcriptionally to the intron (Macias et al 2008). Concordantly, the *RPL30* intron is spliced very slowly, with only 24% spliced after 5 minutes as measured by 4tU-seq, however, SMIT reports that *RPL30* is 90% spliced when the polymerase has progressed only 70 nucleotides beyond the 3’SS. This may reflect intron-mediated inhibition either pausing Pol II prior to 3’SS synthesis, or blocking the second step of splicing, leading to accumulation of intermediates that are not detectable by SMIT. Could other intron-regulated transcripts be regulated by similar co-transcriptional mechanisms?

The Dbp2 protein negatively regulates its own production by binding to its intron to inhibit splicing (Barta and Iggo 1995); its transcript is slowly, yet apparently highly co-transcriptionally, spliced. To investigate at what stage in the transcript’s lifetime this regulation might occur, we looked at the SMIT and NET-seq data in more detail (Fig. 2C). The unspliced SMIT reads decline abruptly ~160nt into the intron, a position coinciding with a T-rich sequence that is presumably hard to align and also locally absent from NET-seq reads. Thus the apparently complete 3’SS-proximal splicing is due to the near-complete absence of unspliced reads near the 3’SS in the SMIT data. However, NET-seq reads continue along the intron and into exon 2, albeit at a low density compared to exon 1, consistent with incomplete passage of Pol II through the intron. Meanwhile, Dbp2 functions as a co-transcriptional RNA chaperone (Ma et al., 2013, 2016). We hypothesize that the intron-mediated negative auto-regulation of *DBP2* splicing occurs co-transcriptionally and results in premature termination or cleavage of the nascent transcript.

**Figure 2:**
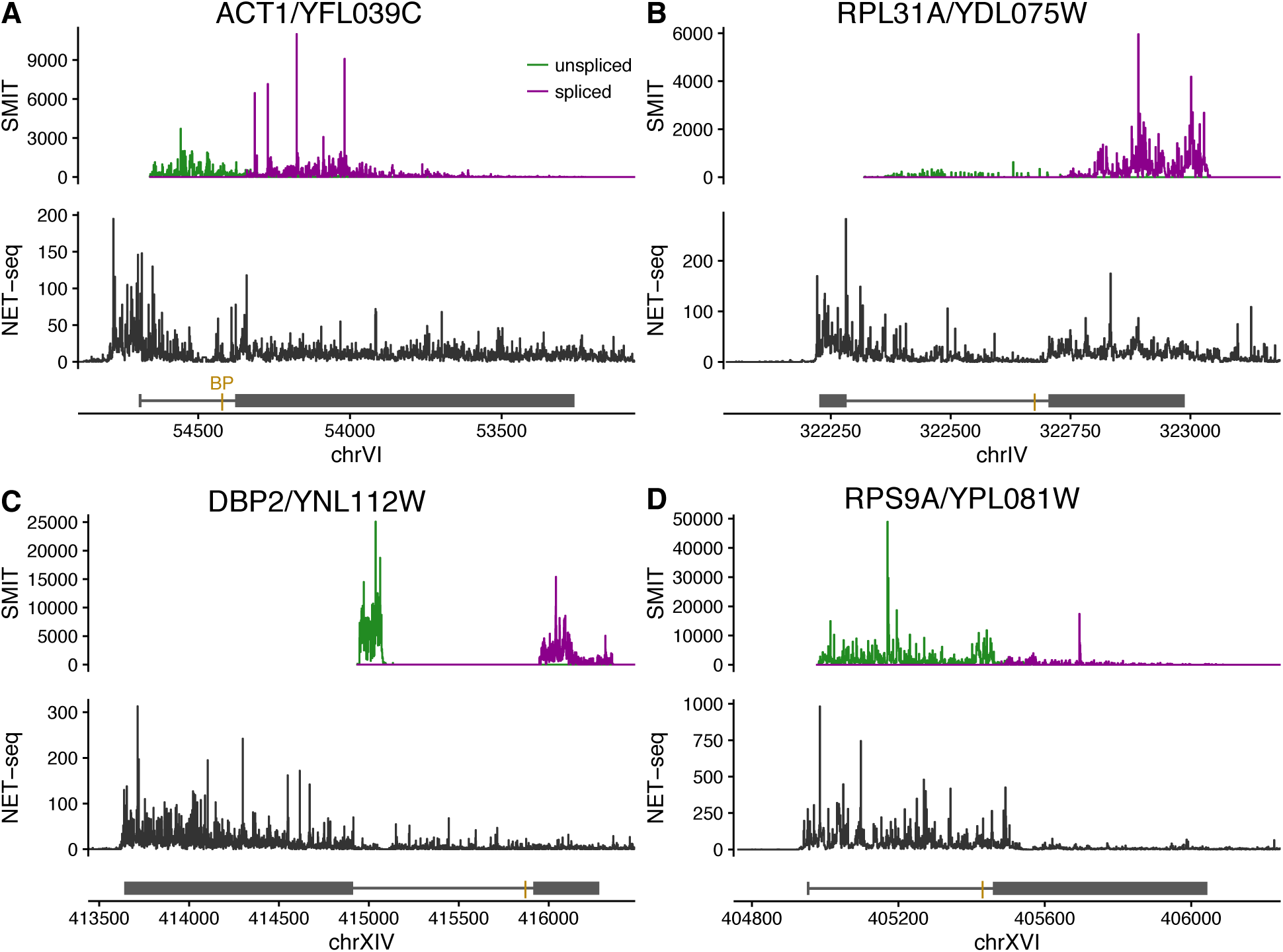
Comparison of SMIT and NET-seq profiles along individual genes, plotted in genomic co-ordinates. Upper panels show 3’ end counts of unspliced (green) and spliced (purple) SMIT reads. Lower panels show 3’ end counts of NET-seq reads. Line diagrams show exons represented by solid blocks, introns by lines, and gold bar represents the branch point (BP); these selected genes are all on the plus strand (5’ left, 3’ right). Note different length scales for each gene. A. *ACT1*/YFL039C is spliced with medium efficiency. B. *RPL31A*/YDL075W is a fast-spliced ribosomal protein transcript: the low ratio of intron:exon 2 reads in SMIT, and the depletion of NET-seq reads towards the 3’ end of the intron are typical of RP transcripts. *DBP2*/YNL112W (C) and *RPS9A*/YPL081W (D) have anomalous profiles.

Likewise, Rps9B protein binds to the *RPS9A* intron and 3’ UTR in chromatin (suggesting that this happens co-transcriptionally) and represses splicing of the *RPS9A* intron (Petibon et al., 2016). Nevertheless the *RPS9A* intron is highly co-transcriptionally spliced (according to SMIT) but with low efficiency (according to nascent RNA-seq). Again, both SMIT and NET-seq show a remarkably low density of reads in exon 2 compared to the intron (Fig 2D). Petibon et al further showed that repression of *RPS9A* splicing is promoter-independent, thus occurs after transcription initiation. We hypothesize that co-transcriptional repression of *RPS9A* splicing by Rps9b protein may result in Pol II pausing or premature termination.

These examples illustrate the rich information that can be obtained by comparing datasets from these different approaches, beyond a simple analysis of splicing efficiency, speed or position. Indeed, comparing the SMIT profiles on individual genes highlights again the distinction between RPs and other intron-containing genes, with the ratio of 3’ ends from intron to exon 2 being low for RPs (Fig. S1), despite their long introns. Furthermore, such profiles highlight the intronic snoRNAs embedded in *EFB1* and *IMD4* introns.

More generally, these vignettes emphasize that regulation of transcription is coupled to regulation of splicing.

### Splicing affects transcription elongation and Pol II phosphorylation

Our group and others have observed that elongating Pol II can pause while the nascent pre-mRNA is being spliced. Pol II accumulates transiently, in a splicing-dependent manner, near the 3’ splice site on reporter genes in budding yeast (Alexander et al., 2010) and certain splicing defects give rise to transcription defects at introns, suggesting a transcriptional elongation checkpoint during co-transcriptional spliceosome assembly (Chathoth et al. 2014). It remains unclear how widespread this pausing phenomenon is: at which positions, in which genes, in which organisms, and in which conditions, does transcription wait for splicing, and for how long? Does pausing occur both before and after the first catalytic step of splicing?

Evidence from high-throughput datasets is so far mixed. Carrillo-Osterreich et al. (2010) report transcriptional pausing ~250bp downstream of the 3’SS using tiling micro-arrays of a chromatin fraction, but Carrillo-Osterreich et al. (2016) report no pausing in sequencing of RNA 3’ ends associated with identically prepared chromatin fraction from a single experiment. The first NET-seq dataset (Churchman & Weissman 2011) did not disentangle RNA 3’ ends that could be caused by Pol II pausing from an abundance of reads at the 3’ ends of excised introns. On the other-hand, Harlen et al., (2016), with higher read-depth, report an excess of NET-seq reads on yeast exons downstream of the 3’SS relative to upstream in the introns, and a peak immediately downstream of the 3’SS, that is interpreted as Pol II pausing. In agreement, NET-seq with human cells shows Pol II accumulation after the 3’SS in a splicing-dependent manner (Mayer et al., 2015; Nojima et al 2015).

Splicing also leads to a change in state of the transcription elongation machinery, notably in the modifications of the Pol II CTD. ChIP-qPCR in yeast (Alexander et al., 2010; Chathoth et al., 2014) and ChIP-seq in humans (Nojima et al., 2015) showed a splicing-dependent phosphorylation of Ser5 in the CTD. Furthermore, ChIP-nexus (a modified, more precise, ChIP-seq; Harlen et al 2016) reported that Ser5 phosphorylation rises rapidly after the 3’SS in yeast. Applying a hidden Markov model to mCRAC data, Milligan et al (2016) also detected systematic differences between overall modification states of Pol II on intron-containing versus intronless yeast transcripts, with changes of phosphorylation state coinciding with positions of splice sites.

Are these effects of splicing on Pol II functionally related? If specific modifications in Pol II are required for efficient transcription through the 2^nd^ exon, then conditions that alter these modifications could lead to transcriptional pausing. It has been suggested (Alexander et al., 2010; Chathoth et al., 2014) that Pol II pausing and phosphorylation could be evidence for splicing-dependent transcriptional checkpoints, implicating quality control of RNA processing in regulating transcription.

Clearly, more extensive and detailed analyses are needed to characterise RNA processing-dependent polymerase behaviour genome-wide. If Pol II pausing happens between the first and second catalytic steps of splicing, detecting the pause via sequencing requires splicing intermediates (specifically, intron-exon lariats) to be quantified simultaneously with the end-products. If a subset of (modified) polymerase pauses transiently for splicing, then the signal is likely to be weak in the pool of total polymerase. If polymerase only pauses on a subset of genes, the signal within metagene analyses will be weak. If polymerase pausing is involved in a regulatory response, or is modulated by environmental stimuli that affect RNA processing, pausing might be rare in some conditions, but common in others.

### Future directions

Collectively, the datasets examined here provide compelling evidence that RP transcripts are spliced both faster and more co-transcriptionally than the average transcript. RP genes produce the largest and most abundant class of spliced transcripts in yeast (Ares et al., 1999), and RP gene expression and splicing are coherently regulated in response to a variety of environmental signals (Pleiss et al., 2007, Bergkessel et al., 2011). The majority of RP introns are required for growth in at least one condition (Parenteau et al., 2011), supporting a regulatory role. RP transcripts are also unusually stable in fast-growth conditions and unusually unstable after glucose starvation (Munchel et al., 2011). Which features of RP genes promote their efficient and co-transcriptional processing? Is it their unusually long (for yeast) introns or their near-consensus GUAUG 5’SS? Is it their stereotypical transcription factors or promoter architecture (Knight et al 2014), or distinctive complement of interacting RNA-binding proteins (Hogan et al 2008)? Could it be connected to reciprocal regulation of ribosomal protein and ribosomal RNA production (Lempiäinen & Shore 2009)? Are there distinct spliceosomal factors that favour splicing of RP transcripts? Unraveling the distinctiveness of RP compared to non-RP gene expression is an important ongoing research focus.

The results discussed here were produced using innovative approaches; how robust will the conclusions seem after further methodological development? Protocols, especially those associated with next-generation sequencing, may need extended optimization to distinguish biological sequence-specific signals from noise and artifacts that arise in both sample preparation and data analysis. Ribosome profiling presents an instructive comparison: the initial analysis for yeast reported that ribosomes do not translate rare codons slowly (Ingolia & Weissman 2009), which was counterintuitive to conclusions from evolutionary biology and biochemistry (Hershberg & Petrov, 2008). Active debate continued over several years as many groups adjusted experimental and analytical protocols, particularly regarding the use of ribosome-stalling drugs. Now, there is clear evidence that ribosome profiling does quantify slower translation at rare codons (Hussman et al 2015, Weinberg et al 2016), in agreement with complementary next-generation sequencing approaches (Pelechano et al., 2016).

Nanopore sequencing of nascent RNA would circumvent the length restrictions of short-read Illumina sequencing; direct RNA sequencing promises to circumvent biases caused by reverse-transcription to cDNA (Garalde et al., 2016). Enrichment of spliced transcripts, analogous to SMIT, might be achievable by real-time selective sequencing on a nanopore sequencer (Loose et al 2016).

The work discussed here has quantified splicing and transcription in the model yeast, *S. cerevisiae*, with unprecedented detail and scope: multiple experimental approaches applied to the same system yield richer insights than any one viewed alone. Do these observations extend to other eukaryotes? For *Schizosaccharomyces pombe*, fast, co-transcriptional splicing was reported by long-read sequencing (Carrillo Osterreich et al., 2016) and suggested by 4tU-seq (Eser et al., 2016). NET-seq in mammals supports co-transcriptional splicing, observing Pol II peaks (interpreted as transcriptional pauses) 3’ to a subset of introns (Nojima et al, 2015) and within spliced exons (Mayer et al., 2015). However, very different efficiencies of co-transcriptional splicing have been reported for Drosophila and mouse (Khodor et al., 2012). A variety of gene features, including intron, exon and overall gene length, intron position, splice sites and other sequence elements, RNA structure and synthesis rate, contribute to differences in splicing kinetics (Khodor et al., 2012; Barrass et al., 2015; Eser et al., 2016). Explaining such differences is not simple and, clearly, much remains to be learned.

**Abbreviations**
5’SS, 3’SS: 5’ and 3’ splice sites.: 
BP: branch point
Pol II: RNA polymerase II
CTD: carboxy-terminal domain of Pol II
4tU-seq: sequencing of 4-thio-Uracil labelled RNA
SMIT: single molecule intron tracking
nts: nucleotides
NET-seq: sequencing of the 3’ ends of polymerase II-associated RNA
mCRAC: crosslinking to modified Pol II and analysis of cDNA
RP: ribosomal protein

## Data and Methods

Data were taken from cited publications, where possible with minimal processing to reflect both the experimental and analysis pipelines used. Nascent RNA-seq data by gene (used in Fig. 1A,C) were a personal communication from K. Harlen, corresponding to Fig. 6D of Harlen et al., (2016); SMIT data are from Carrillo-Osterreich et al. (2016) Table S1, and 4tU-seq data from Barrass et al. (2015), Table S7. SMIT intron:exon ratios in Fig. 1D were calculated by aligning raw reads from GEO (<u>GSE70908</u>; pooled from smit_20genes and smit_extended datasets, excluding short RNA data) to the *S. cerevisiae* genome using STAR (Dobin et al, 2013); SMIT profiles in Figs. 2 and S1 were from the same alignments. Intron lengths in Fig. 1D were taken from Table S8 of Barrass et al (2015). In Fig. 1E, mRNA abundances were taken from Csardi et al (2015b; scer-mrna-protein-absolute-estimate.txt from data package) and U1 occupancy from Table S4 of Harlen et al. (2016). NET-seq profiles in Fig. 2 are from the bedgraph files (WT_NETseq) in GEO: <u>GSE68484</u>. Data were processed using the statistical language R (R core team), and plotted with ggplot2 (Wickham 2009). R scripts to create the figures are available upon request.

## Acknowledgments

We thank members of the Beggs lab, Guido Sanguinetti, Stirling Churchman, and Karla Neugebauer, for discussions and comments on the manuscript. This work was funded by the European Union’s Horizon 2020 research and innovation programme under the Marie Sklodowska-Curie grant agreement No 661179 to EW and Wellcome Trust Awards to JB (104648) and the Wellcome Trust Centre for Cell Biology core grant (092076).

---

**Figure S1:**
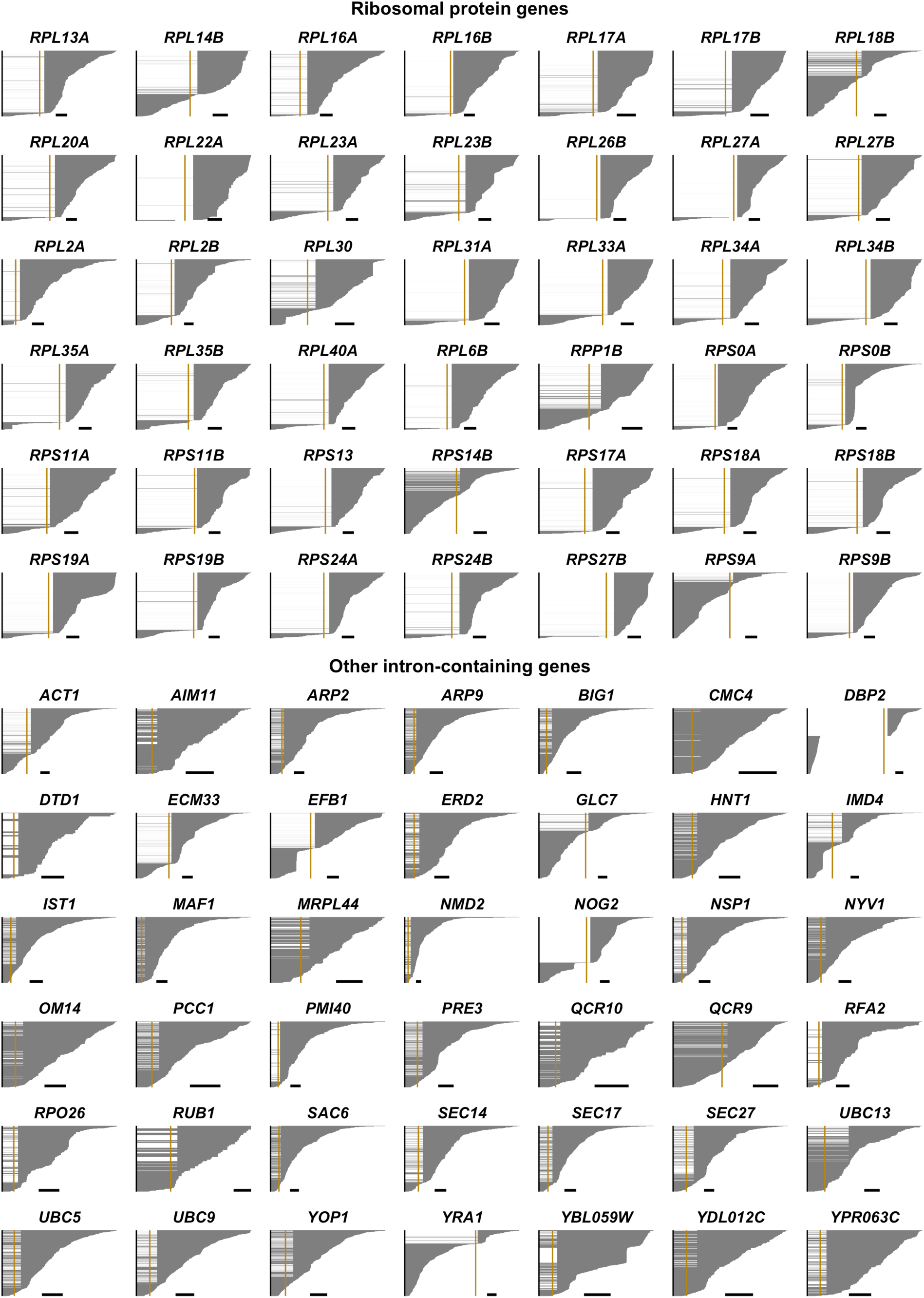
SMIT cumulative profiles for individual genes. Each SMIT read is depicted as a horizontal line from the 5’SS to the 3’ end, excluding the intron for spliced transcripts, and ordered by position of 3’ end. RP transcripts are at the top, and non-RP transcripts below. Note that transcripts are plotted on distinct scales, with 100 nt scale bar for each, and the gold line depicts the BP position. *EFB1* and *IMD4* have an accumulation of reads at the 3’ ends of their intronic snoRNAs.

